# Phosphatidylethanolamine is a phagocytic ligand implicated in the binding and removal of apoptotic and microbial extracellular vesicles

**DOI:** 10.1101/2024.11.18.624161

**Authors:** Ava Kavianpour, Sina Ghasempour, Kirsten J. Meyer, Trieu Le, Ruiqi Cai, Pedro Elias Marques, Justin R. Nodwell, Spencer A. Freeman

## Abstract

The efficient recognition and removal of apoptotic cells by phagocytes is critical to prevent secondary necrosis and maintain tissue homeostasis. Such detection involves receptors and bridging molecules that recognize lipids −normally restricted to the inner leaflet of healthy cells− which become exposed on the surface of dead cells and the vesicles they produce. A majority of studies focus on phosphatidylserine (PS) for which there are well-established receptors that either bind to the lipid directly or indirectly via intermediary proteins. Phosphatidylethanolamine (PE) is even more prevalent than PS in the inner leaflet of mammalian cells and also becomes exposed by the action of scramblases during cell death, though little is known about the effects of PE once scrambled. Here, we report that PE can itself serve as a phagocytic ligand for macrophages by engaging CD300 family receptors. CD300a and CD300b specifically modulated PE-mediated uptake, and this process involved ITAM-containing adaptors and the spleen tyrosine kinase (Syk). For bacteria, which contain PE but largely lack PS in their membranes, we report that PE engagement enabled the binding and uptake of spheroplasts and extracellular vesicles (EVs) that were unsheathed by the cell wall. The inflammatory responses of macrophages to PE particles containing LPS was also curtailed by CD300a expression. Based on these observations, we posit that the direct recognition of PE facilitates mechanisms of clearance that stand to have a broad impact on the immune response.

## Introduction

The lipid bilayers that separate cells from their external environment are made asymmetric in their distribution of lipids by resident lipid-binding proteins [1–4]. Energy-dependent flippases ensure that lipids like PS and PE remain tightly restricted in their incorporation to the cytoplasmic (inner) leaflet while floppases generate enrichment of phosphatidylcholine (PC) and sphingolipids in the exofacial (outer) leaflet [1, 3, 5]. This is a highly conserved process, and though the lipid species concentrations vary in the membrane bilayer between eukaryotes and prokaryotes, the asymmetric nature of plasma membranes is universal [6, 7]. Upon cell death, lipid scramblases of the bilayer become active such that asymmetry is lost [2]. The exposure of lipids normally restrained to the inner leaflet is the major determinant for the removal of the cell by phagocytosis. This has largely been described for PS which has well-established receptors [8–10].

Despite widespread emphasis on PS in initiating the phagocytic programs that remove dead cells, PE is just as prevalent and equally restricted to the inner leaflet of healthy mammalian cells [3]. In healthy cells, like PS, PE is only translocated to the outer leaflet during specific cellular events including at the cleavage furrow when cells divide [11], and to initiate cell-cell fusion that supports the formation of multinucleated osteoclasts [12]. Otherwise, PE is only reported to be scrambled during cell death [13, 14]. While the individual scramblase(s) responsible for exposing PE are not entirely elucidated, the original discovery of the Ca^2+^-activated TMEM16F as a lipid scramblase demonstrated its capacity to scramble PE [15]. The Xkr8 scramblase of the XKR family can also scramble PS and PE in cells undergoing apoptosis [16]. It is noteworthy that in prokaryotes, PE in fact predominates over PS, which is of low abundance [4]. While the lipid membrane of bacteria is masked by surface sugars making up the cell wall, bacteria are known to secrete extracellular vesicles (EVs) devoid of cell wall components [17–19] and their cell wall can also be removed by lysozyme to expose the outer membrane [20]. How host immune cells recognize bacterial debris or EVs is poorly understood.

In that regard, the highly expressed CD300 family of immunoreceptors present in phagocytic cells are known to recognize both PS and PE [21–25]. Here we report that the relative engagement of CD300a, an inhibitory receptor, and CD300b, an activating receptor, dictates the outcome of PE-mediated phagocytosis. PE particles were found to be engaged and phagocytosed by primary macrophages in a manner that was dependent on CD300b. Moreover, PE-receptors function in the binding and uptake of bacterially-derived spheroplasts and extracellular vesicles; CD300a engagement dampens inflammatory cytokine production and release downstream of such particle engagement. Thus, we posit PE as a critical mediator of host-pathogen intracellular communication and an underappreciated phagocytic ligand.

## Results

### Lipid scrambling by the Ca^2+^-activated scramblase TMEM16F exposes exofacial PE

Scrambling of the plasma membrane bilayer is common to all types of cell death. To investigate PS and PE, lipids that are normally kept harbored in the inner leaflet by the activity of flippases (**Fig 1A**), we generated and applied established lipid probes. To this end, PS can be visualized using recombinant Lactadherin C2 fused to GFP (LactC2-GFP) [26], while PE can be visualized using Duramycin-iFluor555 (Dura-555) [14]. To validate the specificity of the probes, lipid-coated beads containing either 5% PS or PE and 95% phosphatidylcholine (PC) were incubated with both probes simultaneously and imaged using fluorescence confocal microscopy. Importantly, LactC2-GFP only labelled the lipid-coated beads that contained PS while Dura-555 only labelled those containing PE (**Fig 1B**). We found that Annexin V behaved similarly to LactC2-GFP and use both probes interchangeably. To demonstrate the exposure of PE and PS upon cell death, we treated RAW264.7 cells with staurosporine (STS) for 3 hours to induce apoptosis. This led to the scrambling of both PS and PE to the outer leaflet of the plasma membrane, as well as the formation of vesicles/blebs (**Fig 1C**).

**Figure 1.**
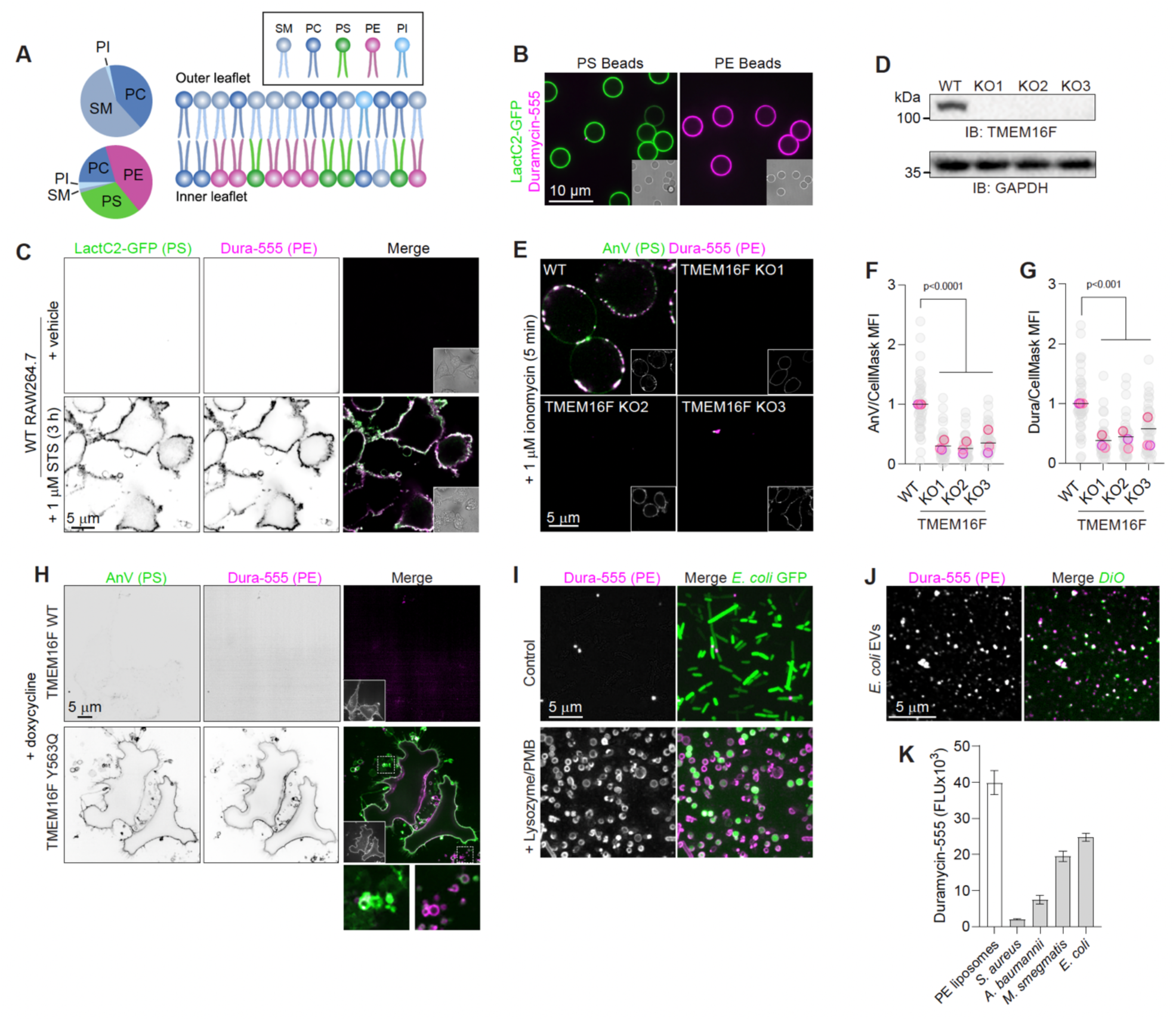
PE is exposed on the surface of apoptotic cells and bacterial vesicles. A) Model based on Doktorova et al., 2023. B) Labelling of lipid-coated beads containing 95% PC and 5% of either PS or PE as indicated with both LactC2-GFP and Dura-555 which were coincubated with the beads for 5 min (n=3). C) WT RAW264.7 cells treated with 1 μM STS to activate cell death pathways and lipid scrambling. Cells imaged live with LactC2-GFP and Dura-555 (n=3). D) TMEM16F KO RAW264.7 cells validated by western blot (n=3). E-G) WT and RAW264.7 TMEM16F KO cells treated with 1 μM ionomycin for 5 min. In E), representative images of cells with AnnexinV and Dura-555 (n=3). The ratio of AnnexinV in F) and Dura-555 in G) to the cell membrane marker, CellMask, are quantified. Grey dots represent fields of view, colored dots represent the mean of the biological replicate. Line represents grand mean. H) HeLa cells induced with doxycycline to express WT or a constitutively active form of TMEM16F (Y563Q). Cells imaged with AnnexinV and Dura-555. Scale bars, 10 μm. I) Control GFP-expressing *E. coli* or those treated with lysozyme and polymyxin B (PMB) stained with Dura-555. Scale bar, 5 μm (n=3). J) Representative image of *E. coli* EVs incubated with Dura-555 and FM4-64. K) Quantification of Dura-555 signal for various indicated particles determined by plate reader. The same number of particles were used in each case.

TMEM16F, a Ca^2+^-activated lipid scramblase, is reported to be essential for the scrambling of PS in response to sustained, elevated concentrations of Ca^2+^ in the cytosol in various cell types [15]. While TMEM16F can scramble a diverse array of lipids including PS, PE, and phosphatidic acid among others, the role of TMEM16F in scrambling PE during programmed cell death has not been established. To establish this, we generated TMEM16F knockouts in RAW264.7 cells (**Fig 1D**) and treated them acutely (5 min) with the Ca^2+^ ionophore, ionomycin, to increase cytosolic [Ca^2+^] and the stimulation of Ca^2+^-activated scramblases. In WT cells, this treatment led to the robust exposure of both PS and PE on the exofacial leaflet of the membrane bilayer as judged by the accumulation of Annexin V (AnV) and Dura-555 on the surface of the cells (**Fig 1E**). This contrasted with TMEM16F KO cells which did not expose PS or PE (**Fig 1E-G**), as previously reported by others in non-macrophage cell types upon Ca^2+^ influx [27, 28].

Importantly, increasing cytosolic Ca^2+^ with ionophores has effects on the plasma membrane beyond lipid scrambling. It also leads to the activation of phospholipases which hydrolyze PI(4,5)P_2_, causing delamination of the cytoskeleton from the membrane, and to the activation of many other pathways that grossly change cell morphology. To determine if the activation of TMEM16F, independent of elevating Ca^2+^, is sufficient to cause scrambling of PE, we generated cell lines that can be induced with doxycycline to express WT TMEM16F- or TMEM16F Y563Q-GFP, a form of the scramblase that is made constitutively active even without increased cytosolic Ca^2+^ [29]. Upon treating the cells with doxycycline, TMEM16F-GFP was synthesized and localized to the secretory pathway and, prominently, to the surface membrane (**Fig 1H**). Convincingly, the expression of the active but not WT form of the scramblase let to the scrambling and exposure of both PS and PE (**Fig 1H**). Interestingly, these lipids could be found to segregate in the scrambled cells, leading to vesicles composed primarily of PS or PE. This suggests that TMEM16F activation alone leads to the exposure of PE together with PS, but that their lateral distribution in the outer leaflet is inhomogeneous. The ability for phagocytes to recognize PE may therefore seem prudent.

### Bacterial vesicles expose PE

In addition to regions and vesicles of dead cells that are enriched for PE, there are several phagocytic targets that contain PE and not PS. Organelles, including mitochondria for example, do not contain PS but have an abundance of PE [30] and if released by necrotic cells would need to be contained by phagocytosis. Bacteria also contain little to no PS in their cell membranes while having a high concentration of PE [6]. Although bacteria are enclosed in cell walls that mask membrane lipids, their shedding of extracellular vesicles (EVs) [17] or the removal of the cell wall by extracellular lysozyme would expose PE, scrambled in the denuded membrane, to receptors.

We therefore endeavored to first investigate the membrane topology of bacteria upon disrupting their outer membrane bilayer with polymyxin. Specifically, GFP-expressing *E coli* were treated with lysozyme and polymyxin B, which lead to the expected formation of spheroplasts (**Fig 1I**) [31]. Convincingly, the spheroplasts exposed PE, as detected with Dura-555 which was only very rarely exposed in control bacteria, curiously seen at the septum that forms during cell division (**Fig 1I**). Bacteria also produce extracellular vesicles: we found that the supernatant fractions of growing bacteria contained EVs as detected by electron microscopy (**S Fig 1A**) and intimated by lipid detection using the FM-4-64 dye (**Fig 1J**). Interestingly, a majority of the FM-4-64 labeled EVs did not react with succinimidyl esters, suggesting low protein abundance in the vesicles (**S Fig 1B**). Using Dura-555, we found that the vesicles were, however, high in PE (**Fig 1J-K**), naturally leading to an investigation of how bacterially derived EVs or PE particles are recognized and removed from tissues/circulation.

### CD300a and CD300b bind PS and PE and modulate phagocytosis

Few receptors bind directly to PS or PE. Of these, TIM-1, TIM-3, TIM-4 and BAI1 are examples, though their surface expression in unstimulated macrophages is restricted [32, 33]. Work on the CD300 family of receptors has identified PS and PE as being directly engaged by CD300a (an inhibitor receptor) and CD300b (a receptor that associates with ITAM-containing adaptors) which otherwise have remarkably similar ectodomains [21, 25]. In RAW264.7 cells we found that CD300a was expressed significantly higher than CD300b, relative to Abt1 (**Fig 2A**), while TIM-4 was undetectable by QPCR (**S Fig 2**). When challenged with beads coated in 5% PE or 5% PS, these cells bound these targets well compared to PC coated beads, but as anticipated, failed to internalize them efficiently (**Fig 2B-C**).

**Figure 2.**
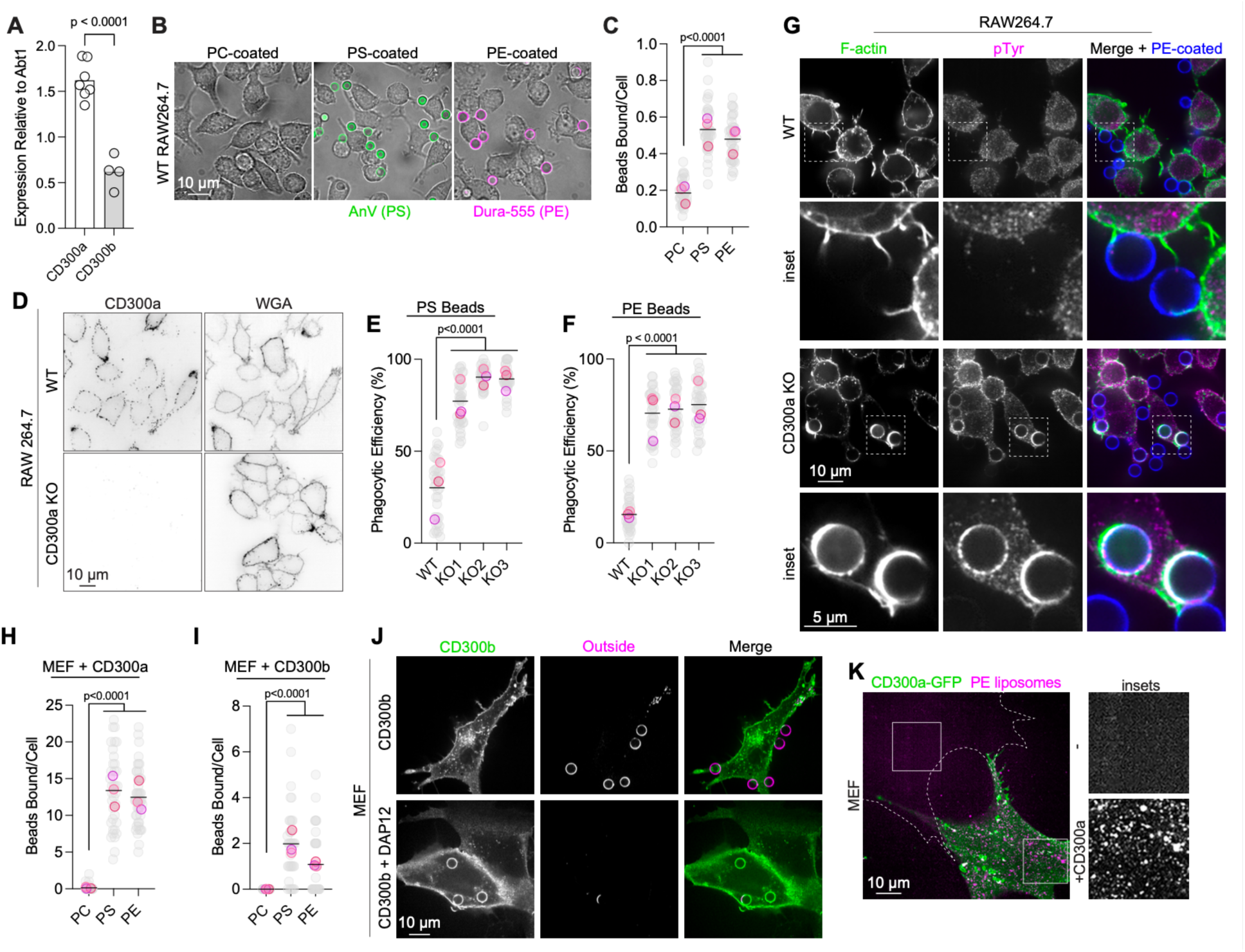
CD300a and CD300b regulate the phagocytosis of PE targets. A) Expression of CD300a and CD300b in RAW cells determined by qPCR, relative to Abt1. Dots represent averages of biological replicates. B) Representative images and C) quantification of PC, PS and PE targets incubated with RAW264.7 cells. Total number of beads bound was divided by total number of cells in field of view (n=3). D) Surface staining of CD300a and wheat germ agglutinin (WGA) of RAW264.7 cells. E-F) Quantification of PS- and PE-coated beads respectively, incubated with WT or CD300a KO cells for 10 min and imaged with E) Lact-C2-GFP (n=3) or F) Dura-Cy5 (n=3). Phagocytic efficiency was calculated by dividing internalized beads by the total beads bound per cell. Data are mean ± SEM, *p* value calculated using Student’s *t* tests (n=3). G) Immunofluorescence of WT and CD300a KO RAW264.7 cells challenged with PE-coated beads for 6 min. H) Quantification of CD300a-OFP expressing and I) CD300b-OFP expressing MEF cells incubated with PS or PE beads in serum free DMEM for 10 min. J) Representative images of MEF cells expressing with CD300b-OFP alone or with DAP12-GFP and incubated with PE-coated beads for 10 min. Cells were imaged live with Dura-Cy5 to label non-internalized beads. Scale bar, 10 μm. K) Representative image of a MEF cell expressing CD300a-OFP incubated with PE liposomes in serum free medium for 10 min. A non-transfected cell in the same field is outlined. In all graphs, grey dots represent fields of view, colored dots represent the mean of biological replicate. Line represents grand mean.

To investigate a putatively inhibitory role for CD300a in PE-particle uptake in RAW264.7 cells, we edited the gene using CRISPR/Cas9. Knockouts were validated using live labelling with an anti-CD300a antibody (**Fig 2D**). We then challenged the WT and KO cells with PE-containing targets, using PS-targets as a comparison. Cells were imaged using LactC2-GFP or Dura-Cy5 which were incubated without permeabilization to mark targets that were bound but not phagocytosed. When compared to unedited (WT) RAW264.7 cells, the CD300a KO cells had a 3- to 5-fold enhanced ability to internalize both PS and PE beads (**Fig 2E-F**). The CD300a KO cells showed no upregulation of either CD300b or TIM-4, suggesting that CD300a functions as an inhibitory receptor that can repress PE-mediated phagocytosis (**S Fig 2**). As CD300a is an inhibitory receptor, capable of recruiting SHP1, SHP2 and SHIP1 phosphatases, we investigated phosphotyrosine signaling and actin polymerization during phagocytosis. WT cells failed to initiate phosphotyrosine signaling at the sites of contact with PE beads and did not advance pseudopodia around the targets, while KO cells were able to generate a complete response (**Fig 2G**). This suggests that CD300a indeed plays a role in inhibiting the phagocytosis of PS and PE targets.

To better study the roles of CD300a and CD300b for PE-mediated phagocytosis in isolation, we opted to express these individually in mouse embryonic fibroblasts (MEF), which are normally non-phagocytic. To this end, CD300a- or CD300b-OFP were ectopically expressed in the MEF cells which were then challenged with PE- or PS-containing targets for comparison. Interestingly, both CD300a and CD300b conferred on cells the ability to bind PE-containing targets to nearly an equal extent as PS-containing targets which was not observed for PC alone (**Fig 2H-I**). CD300a- and CD300b-expressing MEF cells however appeared to only bind but not internalize targets **(Fig 2J)**.

CD300b can associate with the ITAM-containing adaptors DAP10 and DAP12 to promote its signaling. We therefore reperformed these experiments by also expressing DAP12-GFP together with CD300b and imaged with Dura-Cy5 to detect beads not internalized. Here, we found that MEF cells now gained the ability to phagocytose PE beads (**Fig 2J**). The sealed phagosomes that formed in the MEF cells appeared to contain CD300b, further suggesting that the receptor is tightly-engaged to the PE-containing target particles. To investigate the ability of CD300 to bind small targets, e.g. extracellular vesicles, we generated liposomes of <1 μm in diameter that contained 5% PE and 0.5% Cy5-PC. We opted to express CD300a in fibroblasts to limit any internalization and the liposomes were subsequently incubated with the cells. As clearly shown in **Fig 2K**, we found that CD300a conferred on the fibroblasts the ability to bind to the PE-containing liposomes; in contrast, neighboring MEF cells not expressing CD300a failed to bind these liposomes. This would suggest that small lipid particles can be engaged by CD300 family members, as was observed for larger PE-coated targets.

### PE binding in primary macrophages

RAW264.7 cells are poorly efferocytic when compared to primary macrophages. Accordingly, when we employed murine bone marrow derived macrophages (BMDM) as a model, these cells readily internalized PE coated beads. To alter the balance of activating and inhibitory signals, we first overexpressed CD300a-GFP in these cells, leading to a drastic reduction in the phagocytosis of PE-particles (**Fig 3A-B**). Furthermore, blocking CD300b with antibodies significantly reduced but did not eliminate the phagocytic efficiency of the BMDM toward PE targets relative to an isotype control antibody (**Fig 3C**). Many receptors are involved in initiating the signaling cascades that facilitate efferocytosis, however, these tend to converge on the recruitment and activation of Syk. By inhibiting Syk with BAY 61-3606, we found that the uptake of PE targets by BMDM was indeed largely dependent on the kinase (**Fig 3D-E**). These data suggest that there are likely more PE receptors involved in uptake beyond CD300b but that the process largely depends on Syk activity.

**Figure 3.**
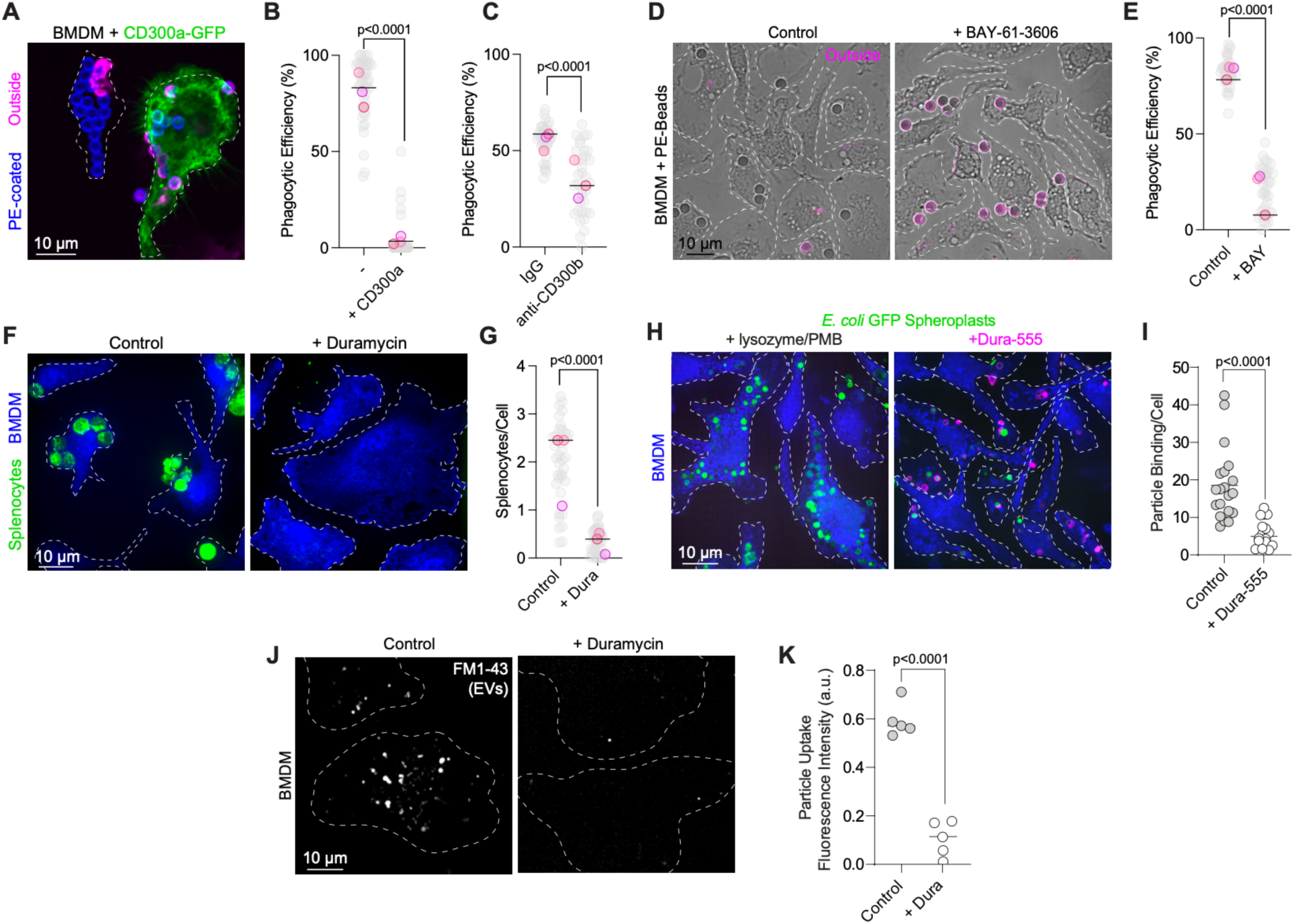
Primary macrophages phagocytose PE targets. A) Representative image and B) quantification of BMDM overexpressing CD300a-GFP challenged with PE-coated beads for 10 min. Dura-555 was used to label beads that were bound but not internalized. C) Quantification of BMDM blocked with IgG or anti-CD300b antibodies and incubated with PE-coated beads for 10 min. Cells were imaged live with Dura-555. D) Representative images and E) quantification of PE-coated beads incubated for 10 min with BMDM treated with vehicle control or 1 μM BAY 61-3606. Cells were imaged live with Dura-555. The phagocytic efficiency was calculated by dividing the number of internalized beads by total beads bound per cell. Scale bar, 10 μm. F) Representative images and G) quantification of CellTracker Blue labelled BMDM (blue) incubated with CellTrace dyed apoptotic splenocytes (green) for 5 min. Apoptosis was induced by incubating splenocytes in 1 μM staurosporine for 3 h. Cells were fixed and imaged. The total number of internalized splenocytes was divided by total number of cells in field (n=3). H-I) BMDM incubated with *E. coli* spheroplasts with or without pre-incubation with Dura-555. Quantification of individual fields of BMDM each containing 5-10 cells. J) Representative images and K) quantification of BMDM incubated with FM1-43 labelled *E. coli* EVs that were either untreated or blocked with Duramycin for 20 min. In all graphs, grey dots represent fields of view, colored dots represent the mean of biological replicate. Line represents grand mean.

To determine if apoptotic cells could also be detected and internalized by their exposure of PE, we first treated primary murine splenocytes with STS for 3-4 h. These targets were determined to be primarily apoptotic but did not become necrotic in this time frame, as determined by their impermeability to propidium iodide (not shown). As expected, once incubated with BMDM, these apoptotic splenocytes were readily bound and engulfed (**Fig 3C-D**). To then block externalized PE, the apoptotic splenocytes were pre-incubated with duramycin, which largely prevented the engagement of targets by BMDM (**Fig 3C-D**). On challenging BMDM with the spheroplasts generated from *E. coli*, we found that binding was also decreased when PE was blocked with duramycin (**Fig 3E-F**). Likewise, we found that the incubation of the bacterial EVs with BMDM resulted in their robust binding and internalization in a manner that seemed to also largely depend on PE since blocking the lipid headgroup with duramycin prevented their uptake (**Fig 3G-H**). Taken together, these data implicate that the ability for macrophages to recognize PE on the surface of phagocytic targets has broad and important effects for internalization.

### Cytokine production and release downstream of PE targets is controlled by CD300 receptors

CD300a shows highest expression in a subset of monocytes, NK cells, and cytotoxic T lymphocytes [34–36]. In mice, the deletion of CD300a increases efferocytosis and inflammation, as orchestrated by monocytes [37]. We noted that in our RAW264.7 CD300a KO cells, there was elevated basal phosphorylation of Syk and Akt relative to WT controls when cultured in DMEM with 5% FBS (**S Fig 3**). All of this suggests that CD300a could modulate the immune response of myeloid cells towards PE-containing targets. Minimal differences were observed between WT and CD300a KO cells in their production and release of cytokines, including when the cells were challenged with PE-containing liposomes (**Fig 4A-L**). However, when the PE liposomes were preincubated with LPS, mimicking targets such as *E. coli* spherocytes and EVs, significant differences were observed in the CD300a KO cells (**Fig 4I-K**). For example, CCL22, also known as MDC, was found to be secreted to significantly higher levels in KO cells relative to WT, suggesting that the PE suppressed secretion when CD300a was present. Interestingly, CCL22 is implicated in rheumatoid arthritis, in which heightened levels were observed in CD300a KO mice [38]. Similar trends were observed for the cytokines LIF and GM-CSF, both of which are capable of heightening inflammation [39, 40]. Other important cytokines released upon LPS stimulation in macrophages such as IL-6, IL-10 and IL-12, did not show significant differences in their release when compared between WT and CD300a KO cells (**Fig 4E-H**). Taken together, these data suggest that PE targets and the PE receptor, CD300a, can have specific effects on inflammation in the context of bacterially-derived ligands.

**Figure 4.**
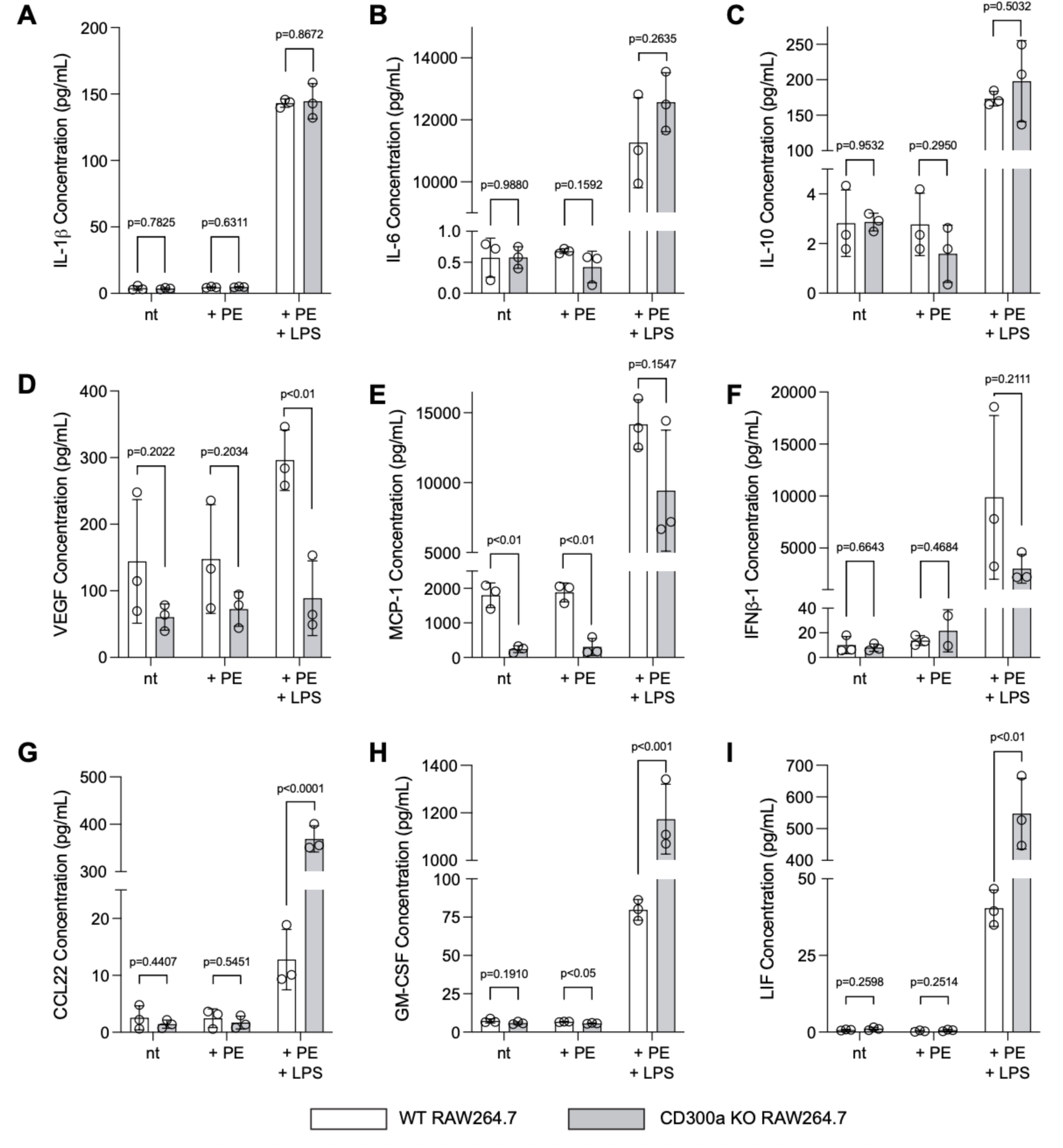
Cytokines released in response to PE and LPS in WT and CD300a KO cells. Cytokines were collected from the supernatants of WT and CD300a KO RAW264.7 cells either untreated or treated with PE liposomes or PE liposomes also containing 100 ng/mL LPS overnight. A) VEGF B) MCP-1 C) IFNβ-1 D) IL-1β E) IL-6 F) IL-10 G) CCL22 H) GMCSF I) LIF. Dots represent biological replicates. Error bars represent SEM. P values calculated using a paired student *t* test.

## Discussion

The collapse of lipid bilayer asymmetry upon cell death is highly conserved. The process is facilitated by the activation of scramblases and the inactivation of energy-supported flippases and floppases. Lipid scrambling is in fact essential for the vesiculation (blebbing) that accompanies apoptosis [41] and membrane repair [27]; we also noted that activation of the TMEM16F scramblase alone was sufficient to cause membrane vesiculation. The outer leaflets of membrane vesicles and debris that are shed from cells thus become rich in lipids normally restricted to the inner leaflet. These same structures tend to exclude the bulky glycocalyx [42] enabling the exposure of scrambled lipid head groups to phagocytic receptors. Surprisingly, we found that PS and PE, once scrambled, could segregate from each other in the plane of the outer leaflet and give rise to vesicles enriched for either lipid. The inherent geometry of PS (cylindrical) versus PE (conical) may help to explain the segregation; the shape of PE may accommodate/generate inward (concave) curvature to enable vesiculation [43, 44]. Given the segregation of the two lipids, it would stand to reason that phagocytic cells must have means to detect either PS or PE.

Most studies have focused on the engagement of PS, either directly or indirectly, since there are select receptors that bind to this lipid and many others that bind to PS-associated bridging molecules like MFG-E8, Gas6, and Protein S [10, 45]. These bridging molecules are engaged by receptor tyrosine kinases of the MERTK family, which rapidly signal for the phagocytosis of the particle [46]. Curiously, MFG-E8, Gas6, and Protein S are all synthesized and released in specific tissues and, in some cases, with a circadian rhythm that supports regular phagocytic programs by resident phagocytes [47]. In certain contexts, bridging molecules may be limiting. However, the direct recognition of PS by TIM-1, -3, and -4 is another important mechanism for initiating the phagocytosis of dead cells [48] which would stand to achieve removal in these cases. There are clearly many inputs and receptors that facilitate the phagocytosis of apoptotic cells and debris which amounts to enough redundancy that other receptor-ligand pairs would not need to be evoked.

Yet, in this report, we find that PE can also serve as phagocytic ligand that leads to phagocytosis that proceeds as efficiently as that supported by the recognition of PS in the absence of serum. While the inner leaflet of the plasma membrane is richly endowed with PS, PE is just as prevalent. Endomembranes, notably the mitochondrial membrane, contain little PS and have high amounts of PE [49]. Chylomicrons may also be much higher in PE compared to PS and PE has been proposed to support removal of lipoproteins from circulation [50]. Additionally, prokaryotic cells, which can shed vesicles devoid of a glycocalyx/cell wall, contain little PS in their membranes [7]. Clearly there are biologically important scenarios where PS is not present to signal the removal of dangerous particulates. In these settings, the recognition of PE may be a critical determinant for phagocytosis. The CD300 family, including CD300a and CD300b, are likely to feature in such situations [21–23, 25].

We found that CD300a is downregulated in inflammatory conditions (**S Fig 2**). The relative expression of these family members across cell types could compartmentalize immune responses. While CD300a does not initiate phagocytosis, it could tether particulates to the plasma membrane and remove these through ongoing membrane turnover, i.e. by macropinocytosis. The inhibitory signaling from CD300a could also be anti-inflammatory [51–53] and may turn off lytic responses of killer cells when they encounter the dead [54]. CD300b, on the other hand, is likely more highly expressed on professional phagocytes, and stands to be a receptor that facilitates the uptake and destruction of PE- and PS-containing particles. Such recognition of PE *and* PS by CD300 family members posits these as aminophospholipid receptors, a category of receptor with critical importance given the prevalence of either aminophospholipid across diverse particle types.

The recognition and removal of bacterial EVs and debris at mucosa is an interesting line of inquiry. For example, it is not yet understood how vesicles produced by the microbiome in the gut, skin, etc. affect immune tolerance or even if these vesicles are bound by resident phagocytes in these tissues. The specific set of receptors that are deployed at this interface therefore remains critical to determine. Moreover, the interactions between microbial species and the use of EVs to impact populations at a distance is also intriguing. Are these vesicles fusogenic and, if so, what mediates the delivery of their payloads between microbes? Curiously, duramycin is an antimicrobial peptide produced by *Streptomyces* [55]. *Streptomyces* protect their own PE from the peptide by its methylation [56]. It is not yet understood precisely how duramycin exerts its effects, but the neutralization of bacterial EVs seems like a possible mode of action. More broadly, the receptors and functions of exofacial PE, whether in eukaryotic or prokaryotic membranes, is important for the field to consider going forward.

## Methods

### Cell culture

RAW264.7 and MEF cells were acquired from the American Type Culture Collection. Cells were cultured in DMEM (Wisent Cat# 319-007-CL) containing 5% FBS (Corning Cat#35-077-CV) and incubated at 37°C with 5% CO_2_. MEF cells were lifted using 0.25% trypsin (Wisent Cat# 325-043-CL). Bone marrow from the long bones from 8–12-week-old C57Bl/6 mice was washed with DMEM with serum and antibiotics then centrifuged. The pellet was resuspended into petri dishes containing 10% L929 fibroblast conditioned DMEM with 10% FBS, penicillin G, streptomycin, and amphotericin B (Wisent Cat# 450-115-EL). Cells were incubated at 37°C for 4-6 days. Bone marrow-derived macrophages (BMDM) were lifted using PBS and 5 mM EDTA, scraped and seeded on coverslips or culture dishes for 24-48 h. Splenocytes were isolated from 8–12-week-old C57Bl/6 mice. The spleen was removed from animals sacrificed by cervical dislocation. The spleen was homogenized by using the flat end of a syringe plunger through a 40 μm cell strainer, collected in a 50 mL centrifuge tube with PBS. Splenocytes were centrifuged at 400 g for 5 min then treated with cold RBC lysis buffer for 5 min. After centrifugation at 400 g for 5 min cells were resuspended in RPMI 1640 medium (Wisent Cat#350-00-CL) with 10% FBS and plated in 6 well plates.

For transient transfections of MEF cells and RAW264.7 macrophages, cells were seeded on glass coverslips within a 12 well plate the day prior. In a 1.5 mL tube, 50 μL serum-free DMEM and 1 μg of plasmid DNA was incubated for 5 min. 3 μL of FuGeneHD (Promega Cat# E2311) was added and incubated for an additional 15 min, then added to the seeded cells. Media was changed 4 h after transfection and cells were imaged the following day. MEF cells were transfected with the following plasmids: CD300a-OFPSpark (Sino Biological Cat# MG50675-ACR), CD300b-OFPSpark (Sino Biological Cat#CG90264-ACG) or CD300b-OFPSpark and DAP12-GFP (Sino Biological Cat#MG53479-ACG). The Invitrogen NeonNxT electroporation system was used to transfect primary BMDM. Differentiated macrophages were lifted and resuspended in R Buffer (Invitrogen A54298-02) and mixed with 1 μg of CD300a-GFP plasmid per well transfecting. Cells were pulsed once at 1400 volts with a width of 30 ms. Electroporated cells were transferred directly to coverslips in 12 well plates.

To generate knockout cells, parental RAW264.7 cells were plated in a 6 well plate. The following day these cells were transfected with a Guaranteed Predesigned CRISPR gRNA plasmid against CD300a (Sigma-Aldrich Cat#MMPD0000125274) or TMEM16F (Sigma-Aldrich Cat#MMPD0000108662). Cells were transfected with 500 μg of DNA and 1.5 μL of FugeneHD per well. Single, live (PI-negative), GFP-positive cells were sorted using FACS into 96 well tissue culture plates. CD300a knockouts were validated using live imaging and qPCR. TMEM16F knockouts were validated by western blot.

### Sleeping Beauty TMEM16F knock-in assay

To test the activity of Ca^2+^ activated scramblase TMEM16F, HeLa cells with doxycycline inducible expression of WT or active mutant Y563Q TMEM16F generated in lab were used. Cells were treated with doxycycline hyclate (Sigma-Aldrich Cat#D9891) for 24 h prior to imaging. Cells were imaged using AnV-647 (AAT Bioquest Cat#20074) and Dura-555.

### Generation of lipid targets

Lipid targets were generated according to protocol by Joffe et. al [57]. Briefly, 4.98 μm silica beads were washed using piranha solution. Small unilamellar vesicles (SUVs) were generated by combining PC (Avanti Polar Lipids Cat#850457C25MG), PC-Cy5 (Millipore Sigma, Cat#850483C), PS (Millipore Sigma Cat# 840035C), or PE (Millipore Sigma Cat#850725C) according to Table 1. To generate targets, 160 μL MOPS buffer, 40 μL SUVs and 20 μL clean beads were incubated for 30 min, centrifuged and gently resuspended in 400 μL of PBS.

**Table 1.** SUV composition.

| Vesicle type | Composition |
| --- | --- |
| PC only | 100% PC |
| PS only | 95% PC, 5% PS |
| PE only | 95% PC, 5% PE |
| Fluorescent PC | 99.6% PC, 0.4% PC-Cy5 |
| Fluorescent PS | 94.6% PC, 0.4% PC-Cy5, 5% PS |
| Fluorescent PE | 94.6% PC, 0.4% PC-Cy5, 5% PE |

### Lipid-binding probes

To detect PS, recombinant LactC2-GFP generated in lab according to protocol by Yeung et al[58] or AnV-647 were used. To detect PE, Duramycin (Dura)-Cy5 (Molecular Targeting Technologies Cat#D-1002) or Dura-555 was generated in lab. Duramycin (Santa Cruz Cat#1391-36-2) and iFluor® 555 succinimidyl ester (Cedarlane Cat#1023) were added to 0.05 M borate buffer, vortexed, and incubated shaking at 500 rpm for 1 h at room temperature. Dura-555 was dialyzed in 1 L PBS to remove unincorporated dye using 2000 Da Slide-A-Lyzer™ Dialysis Cassettes (Thermo Fisher Cat#66205). Collected Duramycin-555 was diluted 1:5 in PBS and stored in the freezer. To validate probes, PS- or PE-targets were incubated with both LactC2-GFP and Dura-555 and imaged. LactC2-GFP was used at 1:50, AnnexinV at 1:500, Dura-Cy5 at 1:50, and Dura-555 at 1:100. Samples with Dura-Cy5 were washed 1x with PBS before imaging.

### Western blot

RAW264.7 cells were plated in 6-well plates. After 24 h, cells were lysed with RIPA buffer (Sigma-Aldrich Cat#R0278) supplemented with protease inhibitors (Thermo Scientific Cat#A32955) and phosphatase inhibitors (Thermo Scientific Cat#A32957). Lysates were transferred to 1.5 mL tubes and spun down at 14000 rpm for 10 minutes at 4°C. Supernatants were collected and protein concentration was determined by Bradford assay (Bio-Rad Cat#5000006). Samples were boiled at 95°C for 5 minutes and run on 4-20% gradient precast gels (Bio-Rad Cat#4561095). Samples were transferred to PVDF membranes, blocked with 2.5% BSA in TBST. Membranes were probed with α-TMEM16F (Sigma-Aldrich Cat# HPA038958), α-GAPDH (AbClonal Cat#AC033), α-SYK (Cell Signaling, Cat#13198), α-phopsho-AKT S473 (Cell Signaling Cat#9271) and α-phopsho-SYK Y525/Y526 (Cell Signaling Cat# 2710) antibodies, detected by HRP-linked secondary antibodies (Cell Signaling Cat#7074,7076). Blots were imaged using Bio-Rad Gel Doc system using ECL (Thermo Scientific Cat# 32106) or Immobilon Forte HRP substrate (Millipore Cat# WBLUF0500).

### Quantitative RT-PCR

RAW264.7 cells were plated in 6-well plates. After 24 h, cells were scraped, and RNA was extracted using E.Z.N.A. HP Total RNA Isolation Kit (VWR CA101414-852). 1 μg RNA from each condition was used to generate cDNA with qScript cDNA SuperMix (VWR Cat#101414-104) according to manufacturing protocols. Into each well in a 96-well plate, 2 μL of cDNA, 5 μL of TaqMan Fast Advanced Master Mix (Thermo Scientific Cat#4444556), 2 μL of water, 0.5 μL of Taqman probe against the and 0.5 μL reference gene *Abt1* probe (Thermo Scientific Cat#Mm00803824) was added. Targets include target *CD300a* (Thermo Scientific Cat#Mm00468054), *CD300lb* (Thermo Scientific Cat#Mm01701741) and *Timd4* (Thermo Scientific Cat#Mm00724713). RT-qPCR was performed using QuantStudio 3 RT-qPCR System (Applied Biosystems, Thermo Fisher Scientific). 3 biological replicates were used per condition. The ΔCt method was used to determine relative gene expression.

### Immunofluorescence

For immunofluorescence staining, PE targets were added to BMDM, centrifuged and immediately fixed with 4% PFA (Electron Microscopy Sciences Cat#15710) for 10 min. Samples were permeabilized with 0.5% triton for 5 min then blocked for 1 h with 5% BSA in PBS. Primary antibodies against phosphotyrosine (EMD Millipore Cat#05-321) was diluted 1:250 in blocking buffer and added to coverslips overnight at 4°C. Samples were washed 3 times with PBS. Secondary antibody donkey anti mouse-488 (Jackson ImmunoResearch Cat#715-545-150) and phalloidin-555 (Cytoskeleton Cat#PHDH1-A) was incubated with coverslips for 45 min and washed 3 times with PBS before imaging.

### Cell death assays

RAW264.7 cells were incubated with 1 μM staurosporine (Sigma-Aldrich S5921) for 3 h or 1 μM ionomycin (Sigma-Aldrich Cat#407950) for 5 min prior to imaging. RAW264.7 TMEM16F KO cells were incubated with ionomycin for 5 min prior to imaging. Cells were imaged with LactC2-GFP or AnnexinV and Dura-555. To quantify PE and PS exposure, fluorescence intensity of AnnexinV and Dura-555 were measured at the plasma membrane, marked by CellMask (Invitrogen Cat#C37608), using Volocity.

### Phagocytosis

For binding and internalization assays with PS and PE targets, RAW264.7 cells, BMDM or MEF cells were plated on glass coverslips into a 12-well plate. Cells were washed with serum free media prior to addition of PS or PE targets. PS or PE targets were mixed with 1 mL of serum free media and added to cells at approximately a 20:1 ratio. Targets were incubated with cells for 10 min and washed gently before imaging live. For experiments inhibiting Syk in BMDM, cells were treated with 1 μM BAY 61-3606 (Cayman Chemical Company Cat#11423) for 10 min prior to addition of targets. For binding assays, beads bound per cell was calculated from images acquired using 25x objective and at least 10 fields of view using the following equation, 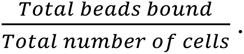 For internalization assays, cells were imaged with LactC2-GFP or Dura-555 to mark outside beads. Phagocytic efficiency was calculated from images acquired using 25x objective, and at least 10 fields of view using the following equation 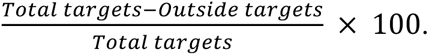

For MEF cell binding assays, MEF cells were transfected with CD300a-OFP or CD300b-OFP. Cells were incubated with PC, PS or PE targets and imaged live. Binding of targets was quantified as described above. For MEF cell internalization assays, cells were transfected with CD300b-OFP and DAP12-GFP. Cells were incubated with PE-targets and imaged live with Dura-Cy5.

### Lipid blocking assay

To generate apoptotic splenocytes, cells were first labelled with CellTrace FarRed DDAO-SE (Invitrogen Cat#C34553) in PBS for 30 min. Cells were centrifuged at 400 g for 5 min and resuspended in PBS then treated with 1 μM staurosporine for 3 h. After treatment, apoptotic splenocytes were centrifuged and resuspended in PBS. To block PE, apoptotic splenocytes were incubated with Duramycin for 2 min, centrifuged, and washed 3 times with PBS. BMDM were plated on glass coverslips in 12-well plates the day before. BMDM were labelled with CellTrace Blue (Thermo Scientific Cat#C34574), then washed with serum free media and incubated with apoptotic splenocytes in serum free media for 5 min. Cells were washed with PBS and fixed with 4% PFA. After fixation cells were imaged.

### CD300b antibody blocking

To block CD300b, BMDM were plated on coverslips in a 12-well plate the day before. Coverslips were transferred to a plate on ice and incubated with CD300b antibody (Bio-Techne Cat#MAB2580) diluted 1:50 in cold HBSS (Wisent Cat#311-515-CL) for 3 minutes. The coverslip was transferred to a well with serum free media and incubated with PE-targets for 10 minutes then imaged. To calculate number of splenocytes bound, splenocytes bound per cell was calculated from images acquired using 25x objective and at least 10 fields of view using the following equation 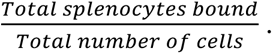

### Live immunofluorescence staining

To label cells for CD300a, cells were seeding on coverslips in a 12-well plate the day before. Coverslips were transferred to a plate on ice and treated with primary antibody against CD300a (Biolegend Cat#147901), diluted 1:50 in HBSS for 3 min. Cells were then incubated in secondary antibody diluted 1:100 in HBSS for 5 min. Coverslips were gently washed 3 times with HBSS and imaged.

### Making spheroplasts and spheroplast phagocytosis

To prepare spheroplasts, 6 ml of overnight culture of GFP *E.coli* was centrifuged at 3100 g, 4°C for 10 min. After the supernatant was removed, the bacterial pellet was used for making spheroplasts [59]. The protocol is modified as follows: after the pellet was washed with 0.8 M sucrose, it was treated with a solution containing 150 µL of 1 M Tris-Cl (pH 7.8), 120 µL of 5 mg/mL lysozyme, and 151 µL of 0.125 M EDTA. Finally, the spheroplasts were collected by centrifugation at 3100 g, 4 °C for 8 min. The supernatant was removed all but 100-150 μl of supernatant left to avoid disturbing the pellet.

The spheroplast pellet was then resuspended in 1 ml of Tyrode buffer (10 mM HEPES, 10 mM D-Glucose, 5 mM KCl, 140 mM NaCl, 1 mM MgCl_2_, 2 mM CaCl_2_). After that, the spheroplasts were treated with 200 μg/ml polymyxin for 13-15 min at room temperature. The suspension of spheroplasts was then divided into 2 equal portions. One portion was added with 1:1000 Dura-555, and incubated at room temperature for 3 min. After that, both samples (with and without Dura-555) were centrifuged at 3100 g, 4°C for 8 min. The pellets were then resuspended with 1 mL Tyrode buffer and added with 100 μg/ml FITC for 5-10 min at room temperature. The samples were then centrifuged at 3100 g, 4°C for 8 min. The pellets were washed with 1 mL Tyrode buffer. The centrifugation and washing steps were repeated three times. The spheroplasts were finally resuspended with 1 mL of serum-free DMEM. BMDMs were plated on coverslips in a 12-well plate the day before. 100 μl of each spheroplast sample was added to CellTrace Far Red DDAO-SE (Invitrogen; Cat# C34553) labeled BMDM. After spheroplasts were added to BMDM, the samples were centrifuged at 1000 rpm for 20 s, then incubated at 37°C, 5% CO_2_ for 10 min. Phagocytosis was done in serum-free DMEM. After that, the samples were rigorously washed with PBS and then fixed with 4% PFA in PBS for 10 min at room temperature. After fixation, cells were imaged.

### Extracellular vesicle isolation, liposomes, and bacterial debris binding assays

Bacterial extracellular vesicles were isolated as described previously [60]. *Escherichia coli* ATCC25922 and *Mycobacterium smegmatis* mc^2^155 were grown in LB, *Staphylococcus aureus* ATCC29213 and *Acinetobacter baumannii* ATCC19606 were grown in Muller Hinton II Broth, at 37°C with shaking at 220 rpm. Overnight cultures from single colonies were diluted; *E. coli* and *A. baumannii* 1:100 and grown for 4 h, *S. aureus* 1:1000 and *M. smegmatis* diluted 1:20 and grown for 24 h. Supernatants from bacteria were centrifuged to pellet cells at 3000g for 10 min, the supernatant was filtered through 0.2µm PES filters, and then concentrated 40x using 10 kDa Amicon Ultra centrifugal filter units (Millipore Cat#UFC901024). Extracellular vesicles were isolated from this concentrated supernatant taking the first 3 mL of elution with PBS after a 4.5 mL void volume of qEV1 35 nm columns (Izon Science Cat.#IC1-35). Transmission electron microscopy images of *E. coli* vesicles negative stained with 2% uranyl acetate were taken on a Talos L120C (TEM imaging & analysis v5.0 SP4). To quantify PE in vesicles, *E. coli* vesicles were incubated with polylysine coated coverslips for 10 min, Dura-555 added and coverslips washed, then FM 4-64 added at 5 µg/mL. To assess PE content between species, vesicle protein concentration was measured by BCA assay (after 30 min incubation with 1% Triton X-100), and vesicles amounting to 500 ng of protein stained with 100 µM Dura-555 for 5 min. Excess stain was then removed by washing 100-fold with PBS using 100 kDa centrifugal filters (Millipore Cat#UFCF10024) at 3000g. Fluorescence of vesicles and PE liposomes stained in the same manner was read in 96-well plates, excitation 500 nm emission 580 nm (Syngery H1, Biotek).

Similarly, to test vesicle binding to macrophages, EVs were stained with FM1-43, the sample was split and half was blocked with Duramycin, then washed with 100 kDa centrifugal filters, and incubated with macrophages for 15 min in serum free media. Cells were imaged live.

### Cytokine analysis

WT and CD300a knockout RAW264.7 cells were seeded in wells of a 6 well plate. Where indicated, cells were treated with either PE liposomes or PE liposomes and 100ng/mL LPS overnight. The following day, supernatant from the wells were collected and spun down at 10000g for 5 minutes to separate cell debris. After centrifugation, the top of the supernatant was collected and frozen at -80°C. Three biological replicates were collected on separate days and sent to Eve Technologies for analysis using the 44-plex discovery assay (Eve Technologies, Cat#MD44).

### Imaging

Images were acquired using a Zeiss Axiovert 200M microscope with a 63x/1.4 NA oil objective or 25x/0.8 NA water objective operated by Volocity software (Perkin-Elmer). Images acquired were analyzed and quantified using Volocity and Fiji (NIH) software.

### Statistical analysis

Data are presented as mean ± SEM. Statistical analysis was performed with GraphPad Prism 8. Statistical significance of data was determined using unpaired Student’s *t* tests with P < 0.05 considered significant.

## Acknowledgments

S.A.F is the recipient of a Canada Research Chair and was supported by PJT-181078 from the Canadian Institutes of Health Research (CIHR) and a Discovery Grant RGPIN-2022-04485 from the Natural Sciences and Engineering Research Council of Canada (NSERC).

## Notes

### Competing Interest Statement

The authors have declared no competing interest.

### Summary of Updates

The manuscript has been revised which resulted in the addition of new data and reordering of the data. Each of the 4 main figures are different from the previous version and now include cytokine analysis from CD300a KO cells and more detailed work on phagocytosis.

