## Supplemental Figures for "Phosphatidylethanolamine is a phagocytic ligand implicated in the binding and removal of apoptotic and microbial extracellular vesicles"

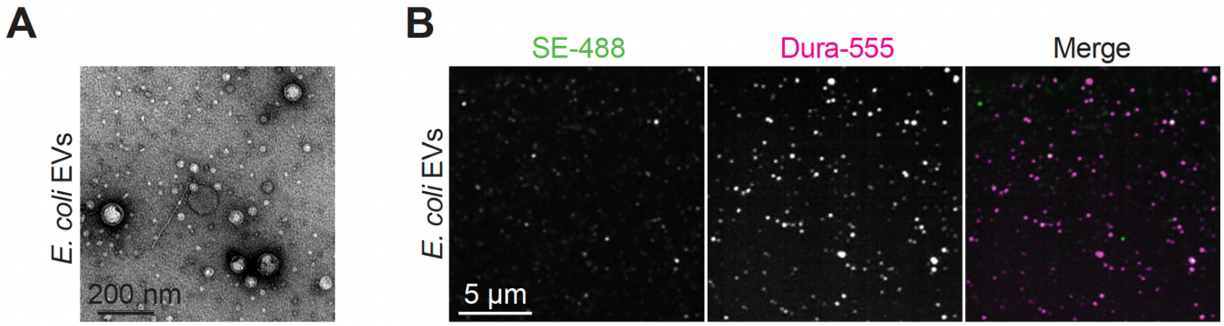


**Supplemental Figure 1 (related to Figure 1).** *Bacterial EVs have limited protein on surface.* A) Representative TEM image of EVs isolated from *E. coli*. B) *E. coli* derived EVs incubated with Duramycin-555 together with Alexa488-succinimidyl ester.

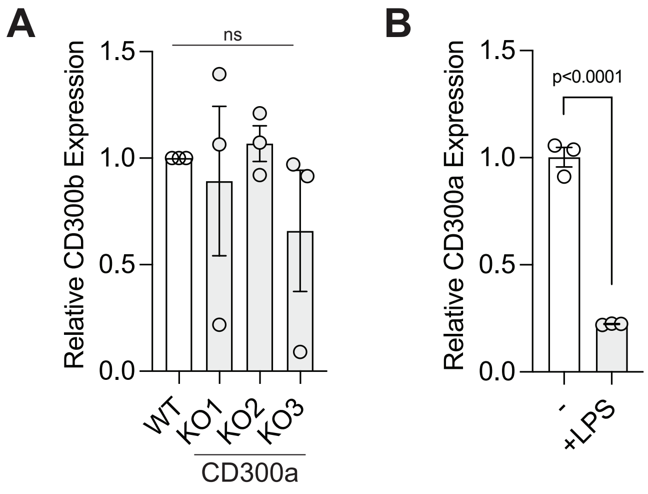


**Supplemental Figure 2 (Related to Figure 2).** *CD300 expression in macrophages.* A) *CD300b* expression in WT and CD300a KO RAW264.7 clones, normalized to *Abt1* expression. Levels of expression of *Timd4* in WT and KO cells was not detected by QPCR. B) Relative *CD300a* expression in WT BMDM either untreated or stimulated with LPS overnight.

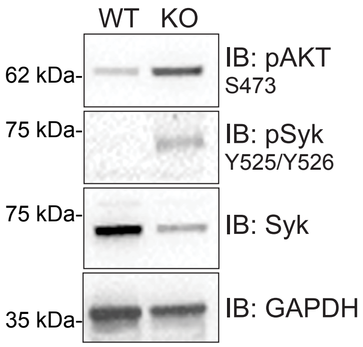


**Supplemental Figure 3 (Related to Figure 4).** *Increased Syk activity in the absence of CD300a.* Western blot of WT and CD300a KO RAW264.7 cells in unstimulated conditions containing complete serum.
